# Spike protein of the SARS-CoV-2 omicron variant interacts with actin

**DOI:** 10.1101/2024.05.16.594608

**Authors:** Ai Fujimoto, Haruki Kawai, Rintaro Kawamura, Akira Kitamura

**Affiliations:** Laboratory of Cellular and Molecular Sciences, Graduate School of Life Science, Hokkaido University, N21W11, Kita-ku, Sapporo, Japan 001-0021; Laboratory of Cellular and Molecular Sciences, Faculty of Advanced Life Science, Hokkaido University, N21W11, Kita-ku, Sapporo, Japan 001-0021; PRIME, Japan Agency for Medical Research and Development, Chiyoda-ku, Tokyo, Japan 100-0004

**Keywords:** COVID-19, Spike protein, actin, fluorescence correlation spectroscopy, protein-protein interaction

## Abstract

The omicron variant of SARS-CoV-2 is responsible for the COVID-19 pandemic, serving as a significant origin for the variants still being detected today. It affects the spike protein that most vaccines used to target when the Omicron strain was discovered. Here, we demonstrate that the receptor binding domain (RBD) of the Omicron variant of SARS-CoV-2 exhibits an increased affinity for human angiotensin-converting enzyme type 2 (hACE2) as a viral cell receptor compared to the prototype RBD. We also identified that β- and γ-actin are Omicron-specific binding partners of RBD. Protein complex predictions suggested that many of the Omicron-specific amino acid substitutions might be involved in the affinity of RBD and actin. Accordingly, we highlight the intriguing observation that proteins expected to localize to different cellular compartments exhibit strong binding.

## 1. Introduction

Coronavirus disease 2019 (COVID-19) is caused by severe acute respiratory syndrome coronavirus-2 (SARS-CoV-2), a positive-strand RNA virus. The worldwide pandemic of this prototype virus (Wuhan-Hu-1) began at the end of 2019 [1]. COVID-19 quickly spread throughout the world, and numerous variants have emerged due to the large number of people infected. Omicron (B.1.1.529), a variant of SARS-CoV-2, was first reported in South Africa in November 2021 [2]. Following the original variant B.1.1.529, several subvariants have emerged (BA.1–5, etc.) [2,3]. Spike glycoproteins (S proteins) on the surface of the virus recognize human cell receptors such as angiotensin-converting enzyme type 2 (hACE2) and mediate membrane fusion between the virus and human cells [4,5]. The S protein is cleaved into the S1 and S2 subunits during viral infection. The S1 subunit contains the receptor-binding domain (RBD) [6,7]. The S protein and the RBD domain itself of SARS-CoV-2 interact with hACE2 with a dissociation constant (*K*_d_) of several tens of nanomolars, and their structural insight has been reported [8,9]. Variant-specific missense mutations are scattered in viral proteins but relatively abundant in the S protein [10]. As the affinity between cell receptors and the virus through its S protein could promote the infection and transmission process, it is important to confirm the variant-specific affinity with ACE2. Although various biochemical and in silico studies have been reported on how these amino acid substitutions affect the affinity of Omicron variants for ACE2 [8,9,11,12], it is not completely conclusive. Fluorescence correlation spectroscopy (FCS) and its advanced method using two-color fluorescence, fluorescence cross-correlation spectroscopy (FCCS), have been used for biomolecular interactions with single-molecule sensitivity [13-15]. The most important feature of FCS/FCCS is a solution measurement system that does not use a solid-liquid interface, as in surface plasmon resonance (SPR). Moreover, since FCS is in fluorescence spectroscopy, it can be used to analyze intermolecular interactions even under cell lysates and in low-purification conditions. Taking advantage of this feature, we compared the interaction strength between the fluorescent protein-tagged recombinant ACE2 and the RBD of both the prototype and the Omicron variants (Wh1 and Omic, respectively) using FCCS.

There has been considerable focus on the affinity of the S protein for cellular surface proteins, the folding process in the ER, and the secretory pathway in producing new viral components and their assembly. However, the interaction of S proteins with intracellular proteins in cellular subcompartments, which are quite distant from typical protein synthesis pathways in cell biology, turns out to be more diverse than initially expected [16,17]. The components of focal adhesion, filopodium, ER membrane, mitochondria membrane, etc. are identified as potential interactors of S protein [16]. These suggest that if S proteins are expressed not only in the secretory pathway but also in the cytoplasm, they can cause various interactome changes that could affect immune responses, cell viability, viral infectivity, and so on.

Here, we demonstrate an enhanced affinity between Omicron RBD and hACE2 compared to the prototype RBD. Additionally, we have identified cytoplasmic actin as a significant interactor of the RBD of the Omicron variant.

## Methods

### Preparation of plasmid DNA

Expression plasmids (pER-mCherry-RBD^Wh1^ and phACE2-eGFP) for protein purification were prepared as previously reported [18]. According to the previous plasmid construction [18], the synthetic oligonucleotides for RBD^Omic^ that carry mutations of B.1.1.529 variant were annealed and inserted into pER-mCherry-C1 (pER-mCherry-RBD^Omic^). For fluorescence imaging, mCherry-tagged ER-sorting signal-lacking RBD plasmids were prepared by vector backbone exchange (pmCherry-RBD^Wh1^ and pmCherry-RBD^Omic^). The plasmid for GFP-β-actin expression was modified as a cDNA-encoding fluorescent tag of YFP-β-actin (TaKaRa-Clontech, Shiga, Japan) was substituted into that encoding eGFP (peGFP-actin).

### Protein purification

Recombinant proteins were expressed in murine neuroblastoma Neuro2a cells (CCL-131, ATCC, Manassas, VA, USA) and purified as previously reported [18].

### Immunofluorescence and confocal microscopy

For confocal imaging, pmCherry-RBD^Wh1^ or pmCherry-RBD^Omic^ (0.15 μg) and p eGFP-actin (0.05 μg) were transfected into HeLa cells (RCB0007, RIKEN BRC, Ibaraki, Japan) using Lipofectamine 2000 (Thermo Fisher) in a cover-glass chamber (5222-004, IWAKI, Shizuoka, Japan) as the manufacturer’s protocol. After incubation for 24 h, the cells were observed using confocal microscopy. For immunofluorescence, HeLa cells expressing mCherry-RBD^Wh1^ or mCherry-RBD^Omic^ were fixed in 4% paraformaldehyde and stained with Phalloidin-iFluor 488 (Cayman, Ann Arbor, MI, USA) according to a previous report [19]. Cell images were acquired using a confocal laser scanning microscope (LSM510 META, Carl Zeiss) with a C-Apochromat 40×/1.2 NA Korr UV-VIS-IR water immersion objective (Carl Zeiss).

### Mass spectroscopic analysis

For mass spectroscopic (MS) analysis, purified proteins were separated using SDS-PAGE, followed by silver staining (423413; Cosmo Bio, Tokyo, Japan). Bands of interest were cut and analyzed using MALDI-TOF-MS, followed by peptide mass fingerprinting that was performed in an outsourced service (Genomine, Pohang, South Korea).

### Western blot

Protein-transferred PVDF membranes (GE Healthcare Life Science) after SDS-PAGE were blocked with 5% skim milk in PBS-T. Antibodies used were anti-GFP HRP-DirecT (#598-7, MBL, Nagano, Japan), anti-mCherry (#Z2496N, TaKaRa, Shiga, Japan) anti β-Actin (#bs-0061R, Bioss, Beijing, China), anti γ-Actin (#11227-1-AP, Proteintech, Rosemont, IL, USA), and a horseradish peroxidase-conjugated anti-rabbit IgG (#111-035-144, Jackson ImmunoResearch, West Grove, PA, USA) antibodies. All primary antibodies were diluted in CanGet Signal Immunoreaction Enhancer Solution 1 (TOYOBO, Osaka, Japan). The secondary antibody was diluted with 5% skim milk in PBS-T. The dilution ratio of all solutions containing primary and secondary antibodies was 1:1000. Chemiluminescent signals were acquired using a ChemiDoc MP imager (Bio-Rad, Hercules, CA, USA).

### Fluorescence cross-correlation spectroscopy (FCCS)

FCCS measurements were performed using a ConfoCor 3 system combined with an LSM 510 META (Carl Zeiss, Jena, Germany) through a C-Apochromat 40×/1.2 NA Korr UV-VIS-IR water immersion objective (Carl Zeiss) as reported previously [18]. Briefly, the relative cross-correlation amplitude (RCA) was calculated as the following equation.

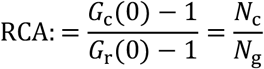

where *G*_r_(0) is the amplitude of the autocorrelation function (ACF) of the mCherry channel at *t* = 0, *G*_c_(0) is the amplitude of the cross-correlation function (CCF) between the eGFP and mCherry channels at *t* = 0, and *N*_g_ and *N*_c_ are the mean number of eGFP-fluorescent and interacting molecules, respectively. Counts per molecule (CPM) were determined by dividing the mean fluorescence intensity by the mean number of fluorescent molecules.

### Prediction of protein complexes

The protein complex of both the Omicron variant and β-actin was predicted using ColabFold v.1.5.5 [20]. The default MSA settings, mmseqs2_uniref_env, were used for prediction, but the AMBER force-field relaxation and templates were not used. The number of recycles was set to 3. The five predicted structures were ranked using pLDDT and pTM scores, and the predicted structure with the highest score was adopted. The positions of amino acids were illustrated using PyMol 2.5.0 (Schrödinger, Inc., New York, NY, USA).

### Statistical analysis

One-way analysis of variance (ANOVA) with post-hoc Tukey’s significant difference test was performed using Origin Pro 2024 (OriginLab Corp., Northampton, MA).

## Results & Discussion

### Increased affinity between RBD of the Omicron variant and hACE2

Purification of the expressed recombinant ER-mCherry-RBD^Wh1^, ER-mCherry-RBDOmic, and hACE2-eGFP expressed in Neuro2a cells was confirmed using western blot for fluorescent tag and silver staining of SDS-PAGE gel (Figure 1, A&B). A single band of purified proteins was observed using western blot (Figure 1A). Although some cellular protein contaminants remained as previously reported [18], the pattern of the bands was similar between ER-mCherry-RBD^Wh1^, ER-mCherry-RBD^Omic^, and hACE2-eGFP (Figure 1B), suggesting that most of them may be nonspecific contaminant proteins. Since, additionally, high purity is not necessarily required for FCCS measurement, the CCFs of the mixture of ER-mCherry-RBD and hACE2-eGFP were measured. The amplitude of the CCF between mCherry-RBD^Omic^ and hACE2-eGFP was higher than that of the CCF between mCherry-RBD^Wh1^ and hACE2-eGFP (Figure 1C). RCA between mCherry-RBD and hACE2-eGFP was significantly positive, and that of Omicron RBD was higher than the prototype RBD (Figure 1D). Thus, RBD carrying the Omicron variant mutation showed a higher affinity for hACE2 than its prototype. Next, RBD is considered to be a monomer, and when the fluorescent brightness of a single molecule (CPM) was evaluated using FCS, the CPM did not change in all samples compared to eGFP and mCherry monomers as controls (Figure 1, E&F), indicating that these samples were monomers even though they interact, and this, moreover, was not aggregates. Therefore, the high RCA of Omicron RBD to hACE2 was not due to the artificial oligomerization of these proteins. Omicron and prototype RBDs share a highly similar binding property to hACE2, but a group of Omicron RBD residues (S496, R498, and Y501) contributes to significant interactions between RBD and hACE2 [11]. Although various in silico and biochemical studies have been reported for the affinity between the Omicron RBD/S protein and ACE2 [8,9,11,12], our FCCS results support the high affinity of Omicron RBD for hACE2.

**Figure 1.**
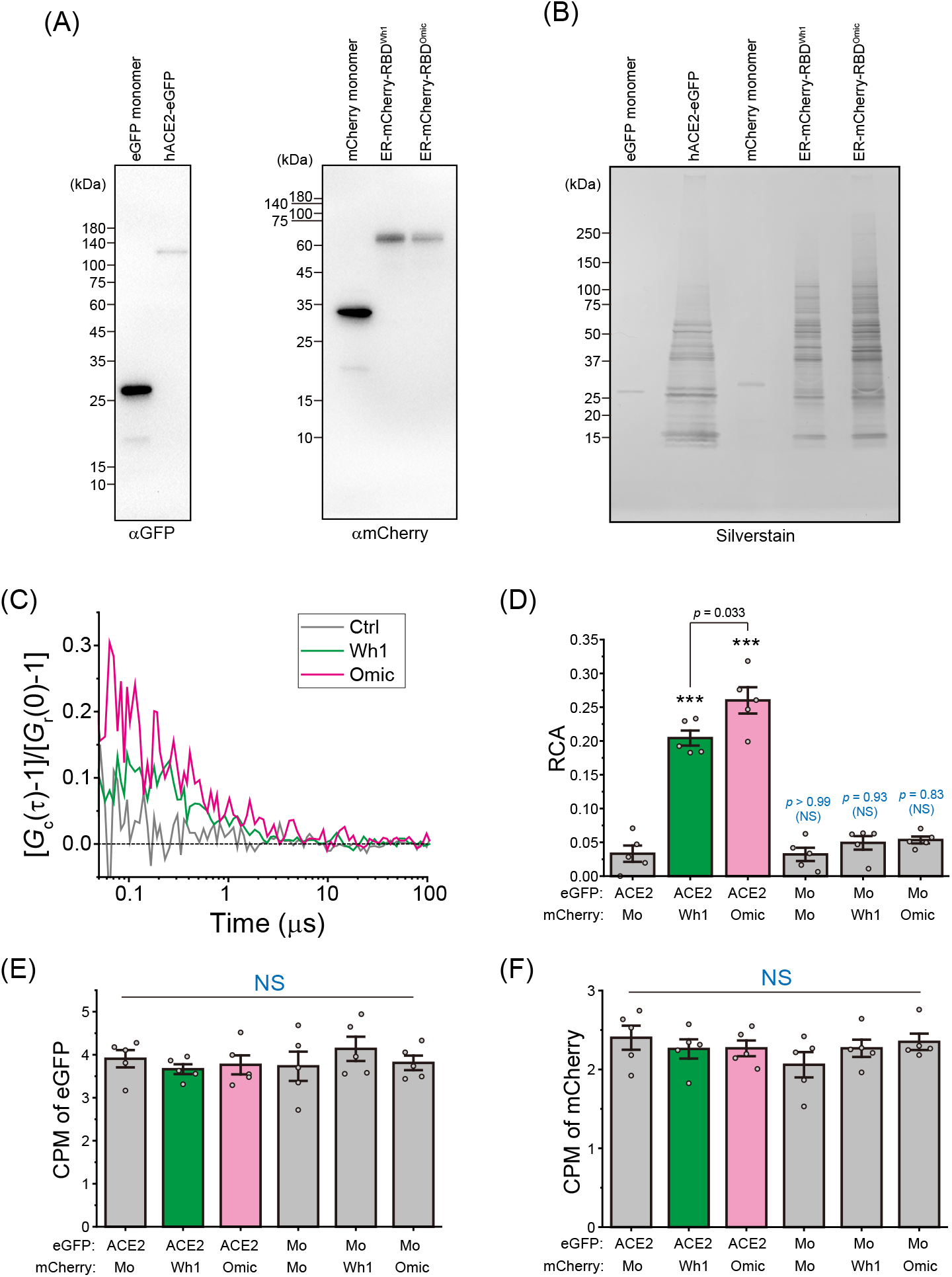
Interaction analysis between ACE2 and RBD of the Omicron/prototype using fluorescence cross-correlation spectroscopy (FCCS). (A) Western blot of purified recombinant eGFP monomer, hACE2-eGFP, mCherry monomer, ER-mCherry-RBD^Wh1^, and ER-mCherry-RBD^Omic^ using anti-GFP and anti-mCherry antibodies (*left* and *right*, respectively). (B) SDS-PAGE gel followed by silver staining of all purified samples represented in (A). The numbers on the left side of the gel image indicate the positions of molecular weight markers. (C) Typical normalized cross-correlation functions ([*G*_c_(τ)-1/*G*_c_(0)-1]) of mixtures of purified mCherry monomers (Ctrl; gray), ER-mCherry-RBD^Wh1^ (Wh1; green), or ER-mCherry-RBD^Omic^ (Omic; magenta) with hACE2-eGFP. The *x*-axis shows the lag time (τ). (D) Relative cross-correlation amplitude (RCA) of indicated two fluorescent color mixtures. (E) Counts per molecule (CPM) of eGFP-tagged proteins during FCCS measurement. (F) Counts per molecule (CPM) of mCherry-tagged proteins during FCCS measurement. (D-F) Mo indicates GFP or mCherry monomers. Bars indicate mean ± SE. Dots indicate independent values. Statistics: \*\*\**p* < 0.001, NS: *p* ≥ 0.05 (not significant).

### Coprecipitation of β- and γ-actin with Omicron RBD

Since FCCS can measure protein-protein interaction with a few small volumes of (~microliters), we did not perform a large-scale cell culture. This is thought to be one of the reasons for not getting highly purified RBDs. However, this is also a condition for coprecipitation of proteins that bind strongly to RBDs, such as immunoprecipitation. From the silver staining pattern of SDS-PAGE of purified RBD proteins for specific binding proteins for the Omicron variant, we found a characteristic ~40 kDa band (Figure 2A). MALDI-TOF-MS analysis revealed that this band contained β- and γ-actin. Western blot using anti-β- and anti-γ-actin antibodies confirmed that the Omicron RBD-specific precipitated band was β - and γ -actin, and was not precipitated with the prototype RBD. (Figure 2, B&C). Since these RBDs labeled with mCherry were expressed in the ER in cells, the binding of cytoplasmic actin to RBDs may occur in the cell lysates.

**Figure 2.**
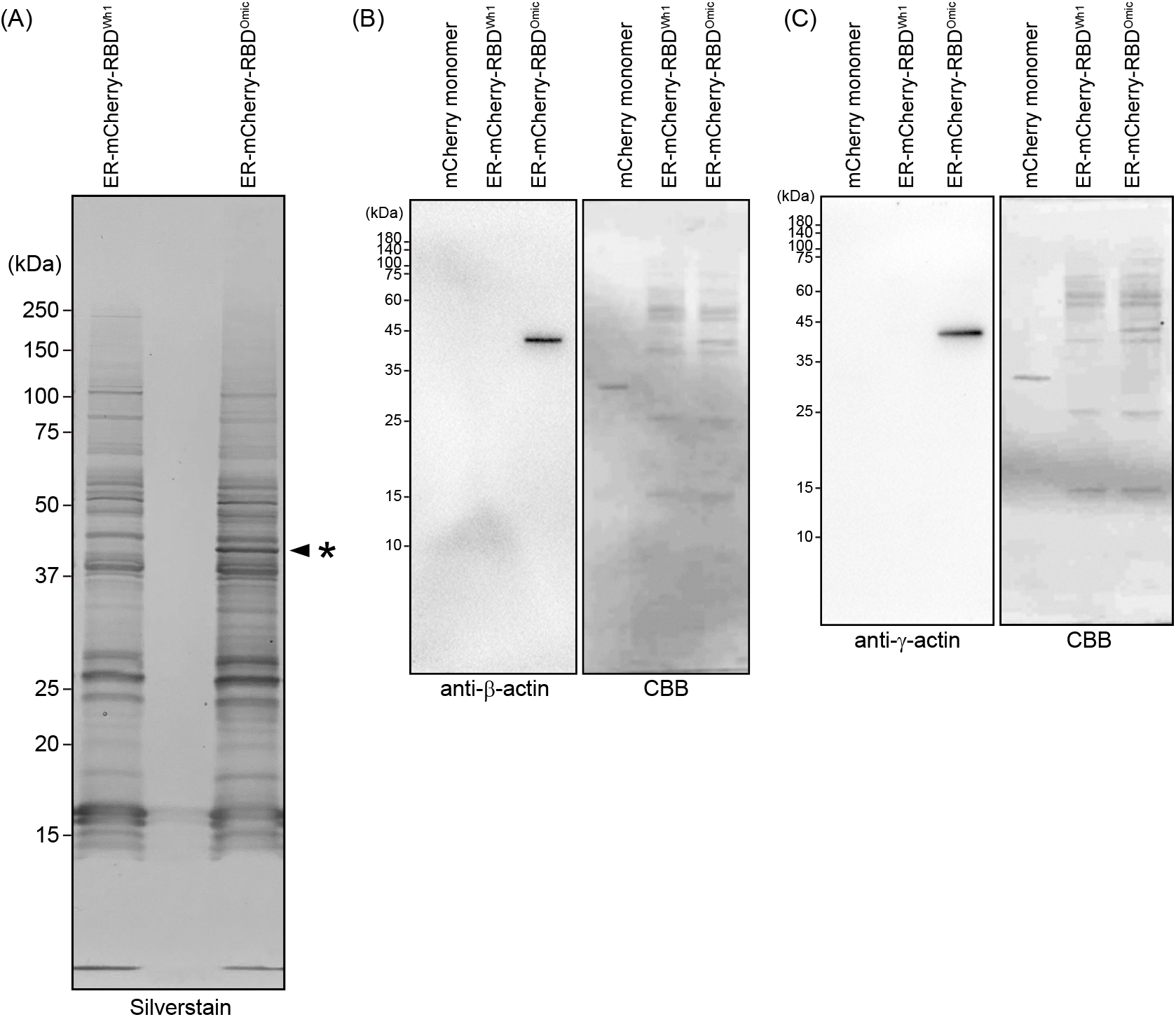
Coprecipitation of beta- and gamma-actin with Omicron RBD. (A) SDS-PAGE gel followed by silver staining of purified ER-mCherry-RBD^Wh1^, and ER-mCherry-RBD^Omic^. The asterisk indicates the band that was specifically coprecipitated with the omicron variant. (B & C) Western blot of the recombinant mCherry monomer, ER-mCherry-RBD^Wh1^, and ER-mCherry-RBD^Omic^ using anti-β-actin and anti-γ-actin antibodies (B and C, respectively). The right images show the membrane stained with Coomassie Brilliant Blue (CBB) after antibody detection.

### Cytoplasmic RBD^Omic^ did not localize on actin filaments

Since actin is a major cytoskeleton, we determined whether cytoplasm-expressed mCherry-RBD binds to actin filaments using confocal fluorescence microscopy. Hereafter, to observe the actin filaments, we used HeLa cells for microscopy. First, actin filaments were visualized by transiently expressed eGFP-β-actin. Although actin filaments attached to the plasma membrane and stress fibers were observed, no colocalization of mCherry-RBD^Omic^ and -RBD^Wh1^ with these actin filaments was observed (Figure 3A). Furthermore, the fluorescent signal of eGFP-β-actin localized in the nonfilament cytoplasmic space and that of mCherry-RBD were not colocalized (Figure 3A). Next, to observe the actin filaments, green fluorescence dye-labeled phalloidin was used for mCherry-RBD-expressing cells; however, no colocalization of mCherry-RBD^Omic^ and -RBD^Wh1^ with these actin filaments was observed (Figure 3B). These results suggest that mCherry-RBD^Omic^ may not bind to actin stationally in the cellular environment.

**Figure 3.**
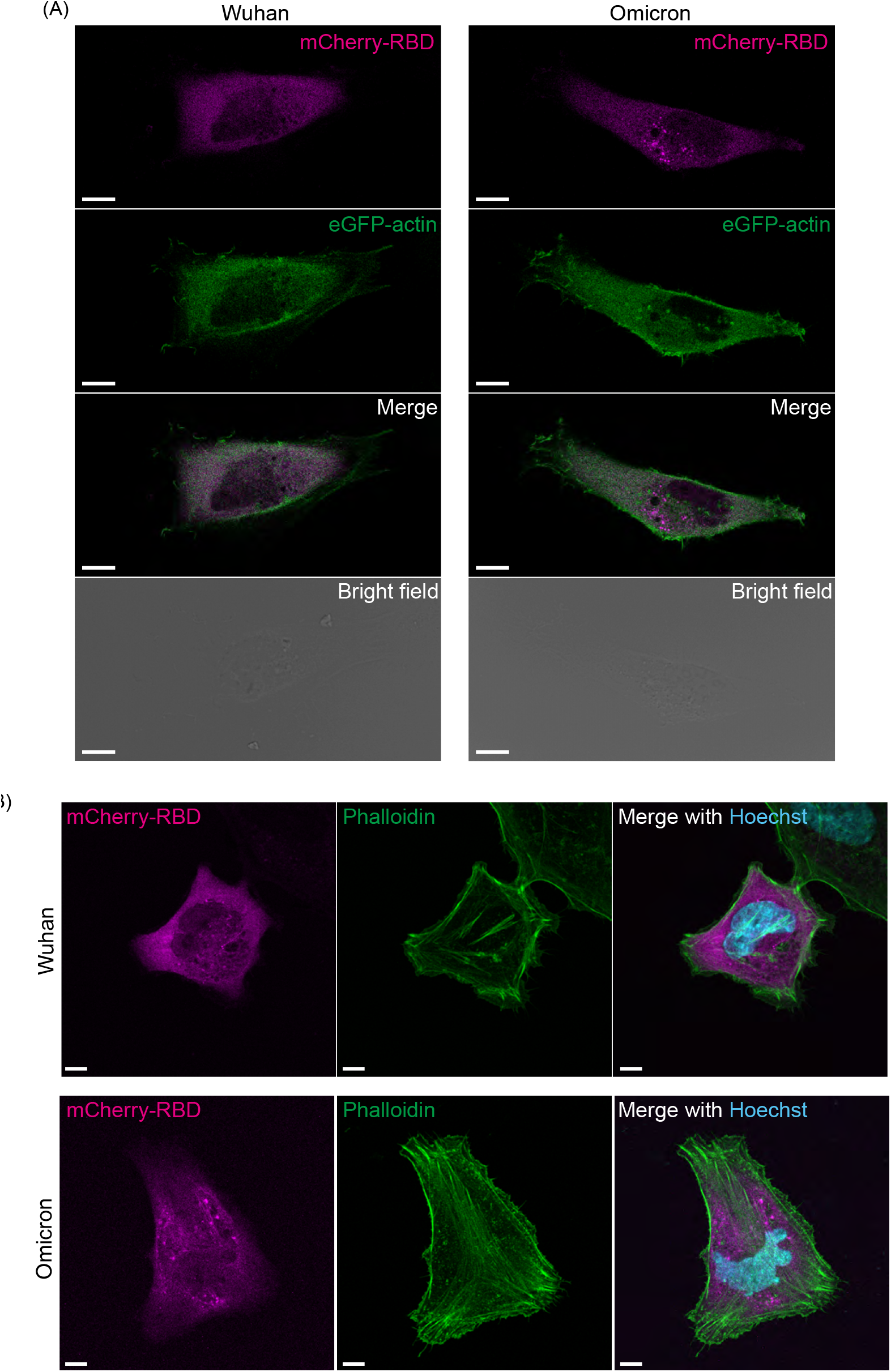
Confocal fluorescent images of actin in HeLa cells expressing mCherry-tagged RBD. (A) Fluorescence images of HeLa cells expressing ER-mCherry-RBD^Wh1^ or ER-mCherry-RBD^Omic^ (mCherry-RBD; magenta) with eGFP-β-actin (eGFP-actin; green). Bar = 10 μm. (B) Fluorescence images of HeLa cells expressing ER-mCherry-RBD^Wh1^ or ER-mCherry-RBD^Omic^ (mCherry-RBD; magenta) stained with Phalloidin-iFluor 488 (Phalloidin; green) and Hoechst 33342 for the nucleus (Hoechst; cyan). Bar = 10 μm.

### The binding of RBD to actin and its physiology

To investigate key amino acids involved in the interaction between RBD^Omic^ and β-actin, we predicted the structure of protein complexes using ColabFold (Figure 4A). Of the total 16 amino acid mutations specific for the Omicron variant compared to the prototype, 9 amino acids (K417N, G446S, T478K, E484A, Q493R, G496S, Q498R, N501Y, and Y505H) clustered within 6 Å from the actin surface (Figure 4B). Among the notable missense mutations in Omicron RBD (N440K, N501Y, and S477N), the N501Y mutation was located 5 Å from the actin surface, while N440K and S477K were located ~15 Å. Among the various missense mutations unique to the Omicron variant, missense mutations that have not been evaluated for binding to ACE2 or viral infectivity might contribute to actin affinity.

**Figure 4.**
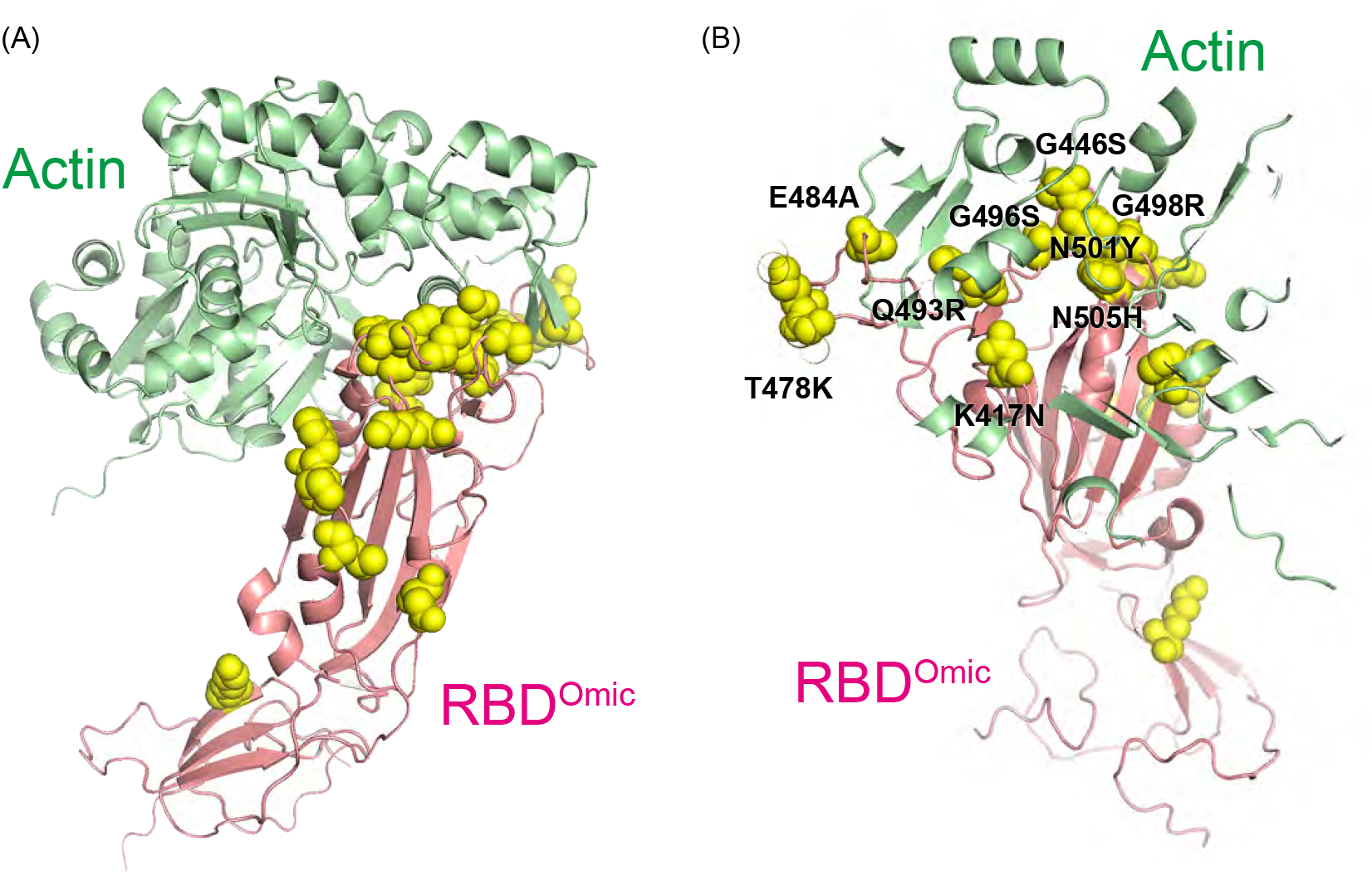
Key amino acids in the predicted complex of RBD and β-actin. (A) Visualization of the predicted protein complex containing both the Omicron RBD (RBD^Omic^; light magenta) and β-actin (light green). Yellow spheres indicate missense mutations in the Omicron RBD. (B) Enlarged view of the complex containing Omicron RBD and actin with increased transparency. Black letters indicate clustered missense amino acids in RBD within 6 Å from the actin surface.

Since S proteins including RBD are translocated into the ER lumen just after its nascent polypeptide chain synthesis, S proteins are unable to meet cytoplasmic actin if this synthetic pathway is followed. However, there is no reason to deny that the nascent polypeptide chain does not leak into the cytoplasm on this pathway, and ORF2 protein of Hepatitis E virus, which undergoes glycosylation in the ER lumen, is retrotranslocated and localized in the cytoplasm, although it is not a substrate of ER-associated protein degradation (ERAD) [21]. Similarly, if the synthesized S protein is leaked into the cytoplasm, it can meet actin. If S proteins may be functionally bound to actin, the possible reason why the colocalization fluorescent signal was not observed in confocal microscopy is that the actin dynamics are extremely fast in living cells due to the involvement of ATP hydrolysis and phosphorylation [22]. But, in cell lysate, the concentration of ATP is diluted, and such dynamics could be stopped. In this situation, S protein and actin remain bound and may be coprecipitated during pulled-down. If this is the case, various regulators of actin dynamics in living cells may be involved in this process.

It remains to be elucidated whether the S protein can be expressed in the cytoplasm in this manner or whether it retrotranslocated into the cytoplasm. If it could be localized in the cytoplasm, whether this binding of S protein would suppress actin dynamics and cellular function, or whether it would be advantageous for virus synthesis, is also a major issue.

### Summary

We have demonstrated through FCCS that the Omicron RBD exhibits an increased affinity for hACE2 compared to the prototype RBD. These findings suggest the feasibility of efficiently detecting interactions between the RBD / S protein and human viral receptors such as hACE2, even amid the emergence of various variants in the future, and that FCCS could serve as a rapid validation method for neutralizing antibodies and small molecule drugs aimed at inhibiting their interaction. Furthermore, we have identified cytoplasmic actin as a significant interactor of the Omicron RBD, while the role of S proteins within the cytoplasm necessitates further elucidation.

## Acknowledgments

We would like to thank Y. Hamada for technical support. The authors acknowledge Open Facility Division, Global Facility Center (GFC), Creative Research Institution, Hokkaido University for the use of a chemiluminescence imager.

## Funding sources

AK was supported by grants from Japan Agency for Medical Research and Development (JP22gm6410028); a Japan Society for Promotion of Science (JSPS) Grant-in-Aid for Transformative Research Areas (A) (24H02286); Grant-in-Aid for Scientific Research on Innovative Areas (22H04826); a grant from Hokkaido University Office for Developing Future Research Leaders (L-Station); a grant from Nakatani Foundation and Hoansha Foundation. AF was supported by the Support for Pioneering Research Initiated by the Next Generation (SPRING) program by the Japan Science and Technology Agency (JST) at Hokkaido University (JPMJSP2119). RK was supported by the Japan Student Services Organization and by the TEIJIN scholarship foundation.

## Author contributions

**Ai Fujimoto**: Writing – review & editing, Methodology, Formal analysis, Data curation, Funding acquisition, Conceptualization. **Haruki Kawai**: Formal analysis, Data curation. **Rintaro Kawamura**: Methodology, Data curation. **Akira Kitamura**: Writing – original draft, Writing – review & editing, Methodology, Formal analysis, Data curation, Supervision, Project administration, Funding acquisition, Conceptualization.

## Conflicts of interest

The authors declare that they have no conflicting interests.

## Notes

### Competing Interest Statement

The authors have declared no competing interest.

